# Speaker Identity is Robustly Encoded in Spatial Patterns of Intracranial EEG for Attention Decoding

**DOI:** 10.64898/2025.12.10.693470

**Authors:** Sukru Samet Dindar, Xilin Jiang, Vishal Choudhari, Stephan Bickel, Ashesh Mehta, Catherine Schevon, Guy M. McKhann, Daniel Friedman, Adeen Flinker, Nima Mesgarani

**Author notes:** Equal Contribution (Sukru Samet Dindar, Xilin Jiang).

## Abstract

The human auditory cortex robustly tracks attended speech, yet it remains unclear if speaker identity is encoded in spatial patterns of neural activity independent of temporal dynamics. Here, we demonstrate that the identity of an attended speaker is reliably reflected in distinct, time-invariant spatial activation maps in human intracranial EEG (iEEG). Leveraging these “neural fingerprints”, we developed a novel framework for Auditory Attention Decoding (AAD) that shifts from traditional temporal envelope tracking to spatial speaker identification. By decoupling the decoding of “who” is speaking from “when” they are speaking, our modular system achieves state-of-the-art speech extraction, particularly in short time windows (<2 seconds) where temporal models typically fail. Furthermore, we observed a reciprocal shift in neural activity during attentional switches, confirming that these spatial codes dynamically track listener intent. These findings establish that speaker identity is a robust, spatially distributed feature in the auditory cortex, offering a high-speed, complementary mechanism for neuro-steered hearing technologies.

## 1. Introduction

The human auditory system possesses a remarkable ability to segregate complex acoustic scenes into distinct auditory objects, a cognitive feat famously known as the “cocktail party” problem (Cherry, 1953). Neurophysiological studies have established that this process involves a dynamic interplay between bottom-up feature encoding and top-down attentional modulation (Ding and Simon, 2012; Mesgarani and Chang, 2012; Patel et al., 2022; Shamma et al., 2011). As a listener focuses on a specific speaker, the neural representation in the auditory cortex selectively enhances the spectrotemporal features of the target voice while suppressing background interference. Translating these neural mechanisms into brain-computer interfaces (BCIs) has given rise to the field of Auditory Attention Decoding (AAD), which aims to steer assistive hearing devices using a listener’s brain activity (Geirnaert et al., 2021; O’Sullivan et al., 2015).

Current AAD approaches, ranging from linear stimulus reconstruction (de Cheveigné et al., 2018; O’Sullivan et al., 2017) to deep neural networks (Ciccarelli et al., 2023; Tanveer et al., 2024) have predominantly exploited a single neural mechanism: the temporal phase-locking of cortical activity to the speech envelope. While robust, this reliance on temporal correlation inherently limits system performance. Because speech envelopes fluctuate relatively slowly, these models require significantly long analysis windows to accumulate sufficient statistical evidence for decoding, resulting in sluggish response times that lag behind natural attentional shifts.

However, the brain utilizes more than just temporal cues to solve the cocktail party problem. Neuroimaging and electrophysiological evidence suggest that the auditory cortex also organizes sound based on time-invariant “what” features, such as pitch, timbre, and speaker identity (Formisano et al., 2008; Norman-Haignere et al., 2015; Walker et al., 2011). Specific neural populations in the superior temporal gyrus (STG) have been shown to tune selectively to these vocal characteristics (O’Sullivan et al., 2019; Patel et al., 2018; van der Heijden et al., 2025). We hypothesized that this feature-based encoding creates a stable, spatial “fingerprint” of the attended speaker that persists independently of the speech envelope’s temporal dynamics. If decodable, this spatial code could offer a complementary, high-speed channel for AAD that bypasses the latency constraints of temporal correlation.

In this work, we introduce a novel paradigm in AAD by targeting the spatial encoding of speaker identity rather than the temporal tracking of speech envelopes. We analyzed intracranial EEG (iEEG) recordings from human participants to determine if the “who” of an attended stream can be decoded solely from spatial activation patterns. Leveraging these neural findings, we developed a modular neuro-steered speech enhancement system that decouples the decoding of speaker identity from the extraction of the speech signal. This approach allows us to train a high-performance speech extractor on massive external audio corpora, guided by the “neural fingerprint” of the target speaker. We demonstrate that this time-invariant, spatially-driven framework outperforms traditional temporal models, particularly in short time windows (<2 seconds), and provides a more robust, biologically grounded solution for neuro-steered hearing.

## 2. Materials and Methods

### 2.1 Participants

The study included 4 participants who were undergoing clinical treatment for epilepsy in one of three medical centers: NYU Langone Health (2 participants), North Shore University Hospital (1 participant), and Columbia University Irving Medical Center (1 participant). Depending on their clinical needs, participants were implanted with either subdural electrocorticography (ECoG) grids and stereo-electroencephalography (sEEG) depth electrodes, or sEEG electrodes alone. All subjects gave their written informed consent to participate, and the research protocols were approved by the Institutional Review Board at NYU Langone Health, North Shore University Hospital, and Columbia University Medical Center.

### 2.2 Experimental Paradigm and Stimuli

Participants took part in an auditory attention task consisting of 28 trials, each with an average duration of 44.2 seconds. In each trial, participants listened to a mixture of two simultaneous and spatially separated conversations presented at equal loudness. Each conversation involved two speakers taking turns, resulting in a total of four unique speakers per trial. The complete dataset featured eight distinct speakers (four female and four male). To simulate a realistic listening environment, background noise (“pedestrian” or “speech babble”) was mixed into the audio at a level of 9 to 12 dB below the conversation streams. Participants were instructed to attend to the conversation that began first, while the competing conversation started with a three-second delay to serve as an attentional cue. The audio was spatialized using head-related transfer functions (HRTFs) and delivered through earphones. To ensure sustained attention, participants performed a behavioral task in which they pressed a button upon detecting intentionally repeated words in the target conversation. All participants successfully completed this task, confirming their engagement. The recorded data was subsequently aligned and segmented into sentences, yielding 280 training, 30 validation, and 50 testing utterances.

### 2.3 Neural Data Acquisition and Preprocessing

Intracranial EEG (iEEG) data was recorded from the implanted electrodes. The raw neural data was bandpass filtered to isolate the low-frequency component between 0.5 and 30 Hz. Electrodes identified as disconnected from the neural tissue were discarded by visual inspection of the neural signals. To maximize brain coverage for the model, the electrodes from all subjects were concatenated into a single dataset.

### 2.4 Proposed Time-Invariant AAD Framework

We designed a novel AAD system centered on decoding time-invariant speaker features from neural signals. The system operates by predicting the cluster label *i* of a fixed-dimensional speaker feature vector *v* ∈ ℝ^*D*^, rather than attempting to reconstruct the high-dimensional vector directly. The model training framework involves a three-step pipeline, as illustrated in Figure 2, which decouples the training of its main components.

**Figure 1:**
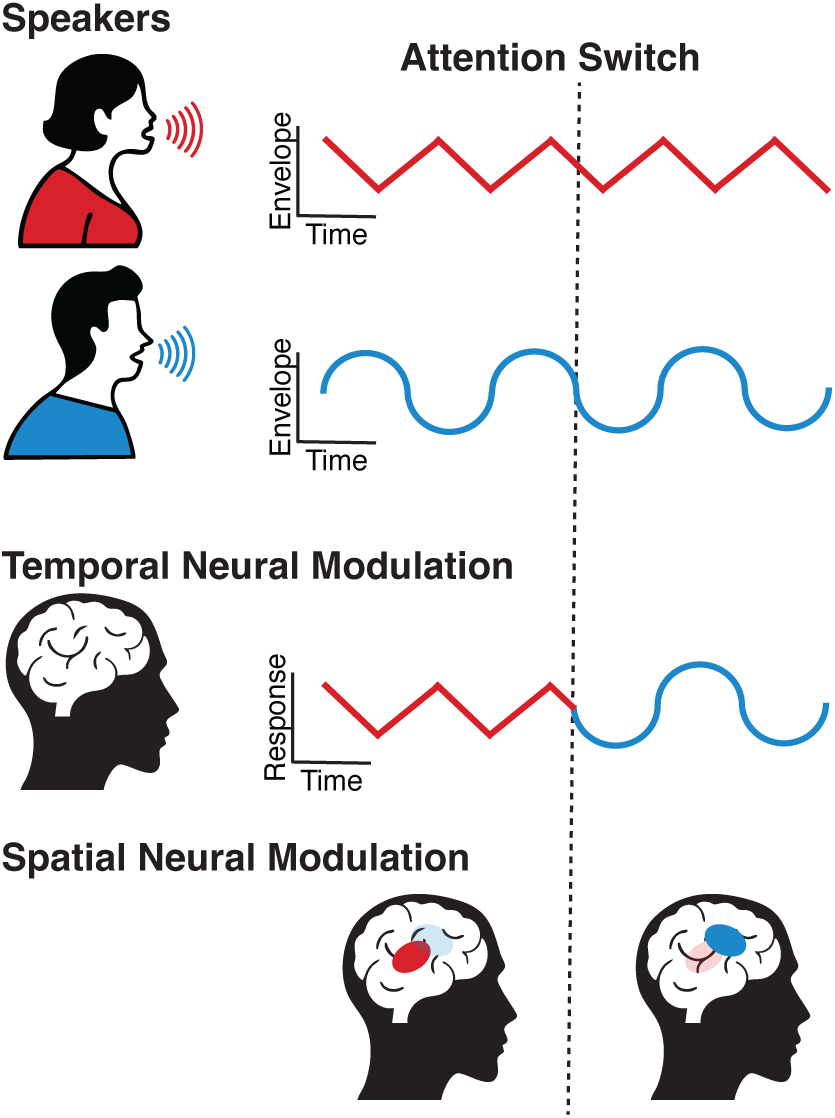
Distinct Neural Signatures of Auditory Attention: Temporal Dynamics versus Spatial Identity. Auditory attention modulates neural activity through two distinct mechanisms. Top: Temporal modulation, where neural activity tracks the time-varying fluctuations of the speech envelope. Bottom: Spatial modulation, where attention selectively engages distinct, speaker-specific neural populations independent of fine-grained temporal timing.

**Figure 2:**
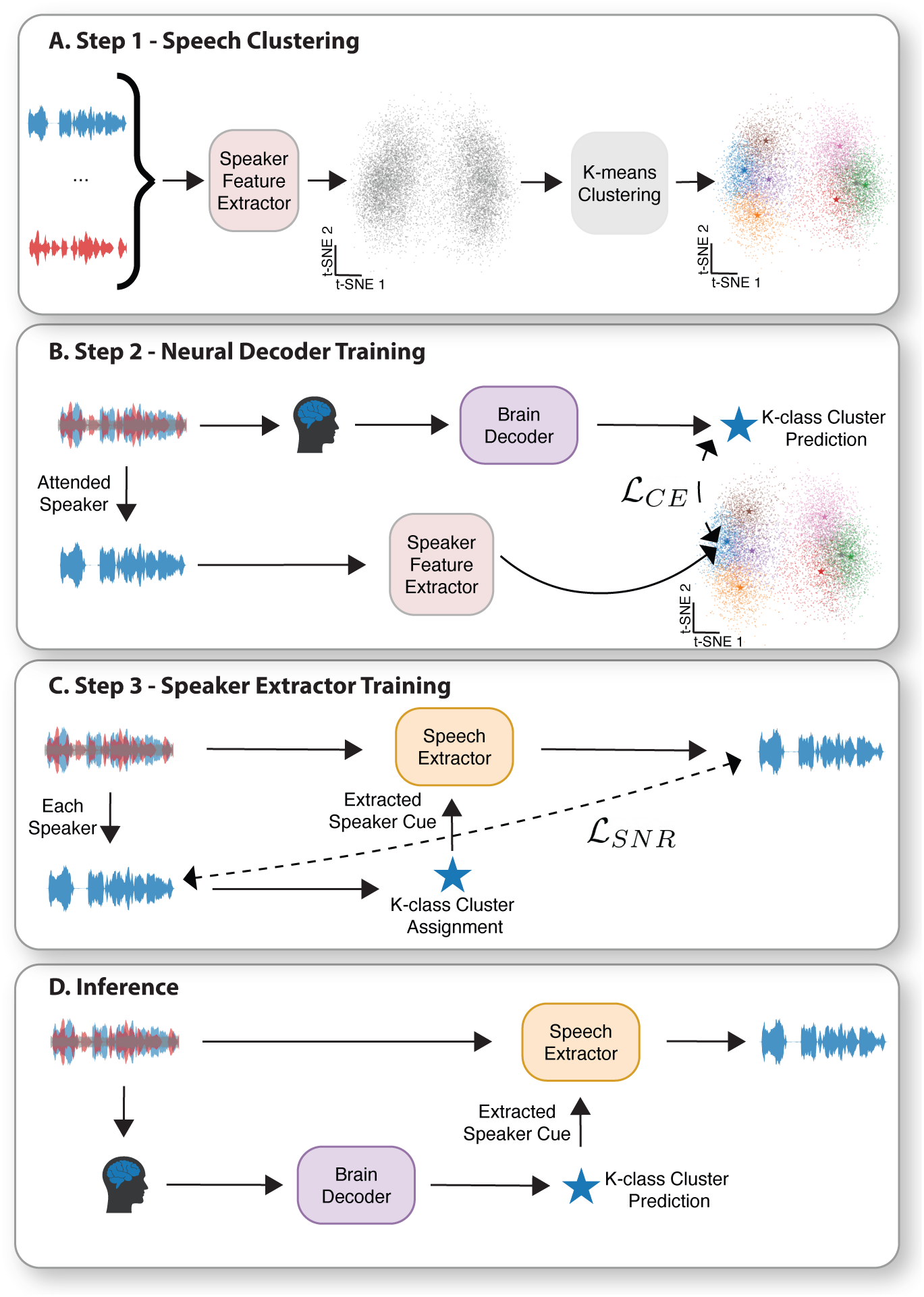
The model operates through three training stages followed by inference. (A) Speaker Feature Clustering: Speaker embeddings from a large external corpus are clustered to obtain time-invariant speaker centroids. (B) Neural Decoder Training: An LSTM-based model is trained to classify the attended speaker’s cluster label directly from iEEG signals. (C) Speech Extractor Training: A speech extraction model is trained independently on speech mixtures to isolate the target voice conditioned on its cluster centroid. (D) Inference: During testing, the brain predictor infers the attended speaker’s cluster label from neural data, and the corresponding centroid guides the extractor to recover the attended speech from the acoustic mixture.

#### 2.4.1 Step 1: Speaker Feature Clustering

To create a general-purpose representation of speaker characteristics, we first performed k-means clustering on speaker features extracted from a large, external speech corpus. We used 10,000 utterances from the LibriTTS dataset (Zen et al., 2019) to fit the clusters, ensuring they were not biased toward the small clinical dataset. Various pre-trained encoders were tested to generate the speaker feature vectors, including a Time Delay Neural Network (TDNN) for speaker verification (x-vectors) (Snyder et al., 2018), StyleTTS 2 (Li et al., 2024), and WavLM (Chen et al., 2022). The number of clusters, *K*, was treated as a hyperparameter. This process yielded a set of cluster centroids {*v*_0_, …, *v*_*K*−1_}. In subsequent steps, any speech segment could be assigned to a cluster *i* by finding the centroid *v*_*i*_ with the minimum ℒ_2_ distance.

#### 2.4.2 Step 2: Neural Decoder Training

With the speaker clusters defined, we trained a neural decoder to classify the attended speaker’s cluster label *i* directly from the iEEG signals. This model was trained on our small, brain-speech paired dataset using a cross-entropy loss function between the predicted label *î* and the ground-truth label *i* derived from the clean attended speech. The predictor’s architecture consists of a bidirectional long short-term memory (LSTM) layer (Hochreiter and Schmidhuber, 1997), a temporal pooling layer, and a final linear layer with a softmax activation function.

#### 2.4.3 Step 3: Speech Extractor Training

The speech extractor module was designed to isolate the voice of the target speaker from a mixed audio signal, conditioned on their speaker cluster centroid *v*_*i*_. Crucially, this component was trained independently on a large-scale, speech-only dataset, using synthetically mixed audio. The model was optimized to maximize the signal-to-noise ratio between the extracted speech *ŝ* and the clean target speech *s*. The architecture is based on Mamba-TasNet (Jiang et al., 2024), which includes a waveform encoder-decoder pair and a stack of bidirectional Mamba layers (Gu and Dao, 2023; Zhu et al., 2024). The speaker cluster centroid *v*_*î*_is used to condition each layer of the network via Feature-wise Linear Modulation (FiLM) (Perez et al., 2017).

#### 2.4.4 Inference

At test time, the system takes a speech mixture and the corresponding neural recording as input. First, the brain predictor processes the neural data to predict the cluster label *î* of the attended speaker. The corresponding centroid *v*_*î*_ is then used as a guiding cue for the speech extractor, which isolates the attended speaker’s voice from the mixture.

### 2.5 Ensemble Model

Recognizing that our time-invariant approach and traditional time-correlation methods might capture complementary information, we developed an ensemble model. This model integrates predictions from our speaker decoding system and a conventional Canonical Correlation Analysis (CCA) based AAD model (Cheveigné et al., 2018). We trained a Random Forest Classifier (Breiman, 2001) as a predictor that makes a final decision based on a set of features from both models. The six features provided to the classifier were: 1) The correlation value from the CCA model for the attended speech stream. 2) The correlation value from the CCA model for the unattended speech stream. 3) The distance between the predicted cluster’s x-vector and the attended speaker’s x-vector. 4) The distance between the predicted cluster’s x-vector and the unattended speaker’s x-vector. 5) The logit score (confidence) of the brain predictor for the chosen cluster. 6) The length of the time window used for the prediction.

### 2.6 Implementation and Training Details

The LSTM network used for the brain predictor had a state dimension of *S* = 64. It was trained for 30 epochs using the Adam optimizer (Kingma and Ba, 2014) with a batch size of 1 and a constant learning rate of 10^−4^. The speech extractor followed the Mamba-TasNet (M) configuration (Jiang et al., 2024) with *R* = 32 layers and a feature dimension of *M* = 256. It was trained for 50 epochs using the Adam optimizer with a batch size of 4 and a decaying learning rate that peaked at 2 ∗ 10^−4^. The Random Forest Classifier was implemented using the scikit-learn library (Pedregosa et al., 2011) in Python with 100 estimators. It was trained and evaluated on a split half of the test set.

## 3. Results

To investigate the neural encoding of speaker identity, we obtained high-density intracranial EEG (iEEG) recordings from four neurosurgical patients implanted with either subdural electrocorticography (ECoG) grids or stereotactic depth electrodes (sEEG), providing broad coverage of the auditory cortex. Participants performed a realistic “cocktail party” task involving 28 trials of concurrent, spatially separated speech streams delivered via head-related transfer functions (HRTFs). The stimuli were constructed from a diverse pool of eight distinct speakers—balanced for gender (four male, four female) to ensure acoustic feature diversity—and mixed with environmental background noise (pedestrian or speech babble) to simulate naturalistic listening conditions. Subjects were instructed to maintain sustained attention on a designated target speaker while ignoring a competing talker, with engagement verified via a behavioral target-detection task. We analyzed the spatial topography of neural responses across these diverse speaker conditions to determine if the attended speaker’s identity could be decoded solely from the spatial distribution of cortical activity, independent of temporal envelope tracking.

### 3.1 Speaker Identity is Reflected in Spatial Neural Patterns

We first sought to validate the core premise of our framework: that the identity of an attended speaker is reflected in distinct, spatial patterns of neural activity. While previous work has identified speaker-selective sites in the auditory cortex (Formisano et al., 2008; Mesgarani and Chang, 2012; O’Sullivan et al., 2019), it remains unclear if these spatial maps are robust enough to track attention dynamically. To investigate this, we monitored electrode groups with preferential responses to specific speakers during an attentional switch. Crucially, we observed a double dissociation: activity in electrodes selective for the first speaker dropped immediately following the shift, while activity in electrodes tuned to the second speaker rose concurrently (Figure 3A).

**Figure 3:**
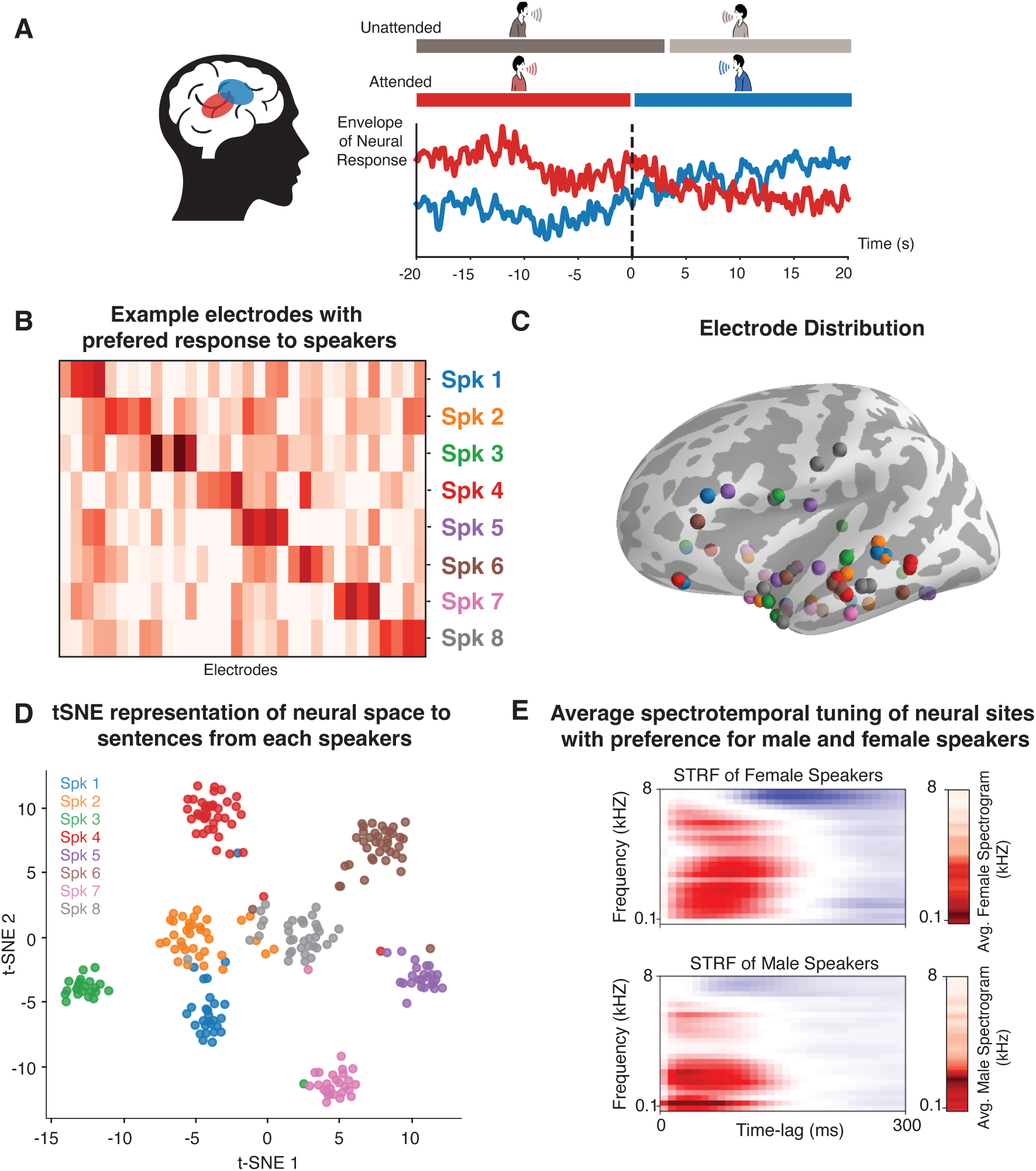
Spatial organization and dynamic tracking of attended speaker identity in the auditory cortex. (A) Average responses of speaker-selective neural sites to two different attended speakers. (B-C) Average neural responses of selected neural sites grouped by the attended speaker, and corresponding electrode distributions visualized on a brain map. (D) t-SNE visualization of LDA-transformed neural responses, highlighting the seperability of neural response patterns with respect to the attended speakers. (E) Spectrotemporal receptive fields (STRFs) and average acoustic spectra reveal distinct tuning properties.

This speaker-specific encoding was evident across the entire cohort of eight speakers. Figure 3B shows the time-averaged responses of selected example electrodes, revealing distinct activation patterns for each attended speaker. These speaker-selective electrodes are distributed spatially across the cortex with no apparent anatomical organization, as visualized in Figure 3C, reflecting the multidimensional acoustic tuning selectively of the auditory cortex (Mesgarani et al., 2008; Walker et al., 2011).

To further visualize the separability of these neural representations, we used Linear Discriminant Analysis (LDA) (Blei et al., 2003) to project the neural responses for each sentence into an eight-dimensional space (one for each speaker) and then used t-SNE (Maaten and Hinton, 2008) for visualization. As shown in Figure 3D, the neural responses form distinct clusters for each speaker, confirming that speaker identity is a decodable feature. Interestingly, these clusters also exhibit a higher-order organization based on gender, with male speakers (1-4) and female speakers (5-8) grouping separately along the horizontal axis.

To relate the observed preferential response of neural sites to speakers to their acoustic tuning, we estimated spectrotemporal receptive fields (STRFs) for electrodes that showed preferential responses to male versus female speakers. STRFs are linear models that reveal the spectrotemporal features to which a neural site is most sensitive (Theunissen et al., 2000). The analysis showed that some neural sites were tuned to frequency ranges characteristic of male voices, while others were tuned to frequencies more typical of female voices (O’Sullivan et al., 2019; Patel et al., 2018). These findings provide a strong neuroscientific basis for a model that decodes attention by mapping spatial patterns of neural activity to time-invariant speaker features.

### 3.2 Attended Speech Extraction Performance

We evaluated our system on five key metrics: signal-to-noise ratio (SNR), scale-invariant signal-to-distortion (SI-SDR) (Le Roux et al., 2019), perceptual evaluation of speech quality (PESQ) (Rix et al., 2001), word error rate (WER) (Radford et al., 2023), and speaker similarity (SIM)(Chen et al., 2022). As shown in Table 1, our proposed model, using x-vectors with *K* = 8 clusters (xvect-8), achieves strong performance across all metrics and substantially outperforms conventional AAD baselines, including the deep learning-based brain-informed speech separation model (BISS) (Ceolini et al., 2020), and systems based on envelope or spectrogram reconstruction. The ensemble model, which combines our time-invariant approach with a traditional CCA-based model, achieves the best performance overall. Notably, the final speech extraction accuracy of our model is higher than the brain predictor’s classification accuracy. This is because even when the predictor selects an incorrect cluster, its proximity in the feature space to the correct cluster is often sufficient to guide the speech extractor to the correct speaker in the mixture.

**Table 1:** Performance comparison of our model to AAD systems with time-varying features.

| Method | SNR | SI-SDR | PESQ | WER% | SIM% |
| --- | --- | --- | --- | --- | --- |
| time-invariant speaker feature decoding (xvect-8) | 12.8 | 10.4 | 2.5 | 14.3 | 94.8 |
| ensemble model (CCA + time-invariant) | **13.0** | **12.7** | **2.6** | **9.0** | **95.5** |
| extraction w/ envelope (BISS) | 6.3 | 4.0 | 1.7 | 37.3 | 88.3 |
| separation + mel spectrogram reconstruction | 12.0 | 9.6 | 2.4 | 15.2 | 94.1 |
| separation + envelope reconstruction | 11.2 | 7.8 | 2.3 | 20.4 | 90.7 |
| separation + mel spectrogram CCA | 12.3 | 10.5 | 2.4 | 14.5 | 94.1 |
| blind separation (without cue) | 6.3 | -8.7 | 1.9 | 55.1 | 83.1 |
| unprocessed | 0.1 | 0.1 | 1.3 | 38.4 | 84.4 |

### 3.3 Analysis of Time Window and Attention Shifts

The duration of the time window is a critical parameter for AAD systems, presenting a trade-off between prediction robustness and temporal resolution. Traditional methods that rely on temporal correlations are particularly sensitive to this parameter, as longer windows provide more stable correlation estimates. While our model is not powered by such correlations, its recurrent neural network (RNN) architecture still benefits from the richer temporal context provided by longer data sequences. We evaluated our speaker decoding model (SPK model), a CCA-based model, and the ensemble model across various window sizes ranging from 0.1 to 8 seconds. Figure 4 shows the mean decoding accuracy across subjects, with error bars denoting the standard error of the mean. As shown in Figure 4A, the CCA model’s performance improves with longer windows, whereas our speaker decoding model achieves higher accuracy at moderate window sizes of 1–2 seconds, which aligns with typical sentence durations. Notably, the composite model consistently outperforms both individual models across all window sizes greater than 0.1 seconds, highlighting the complementary strengths of time-invariant and time-varying decoding strategies.

**Figure 4:**
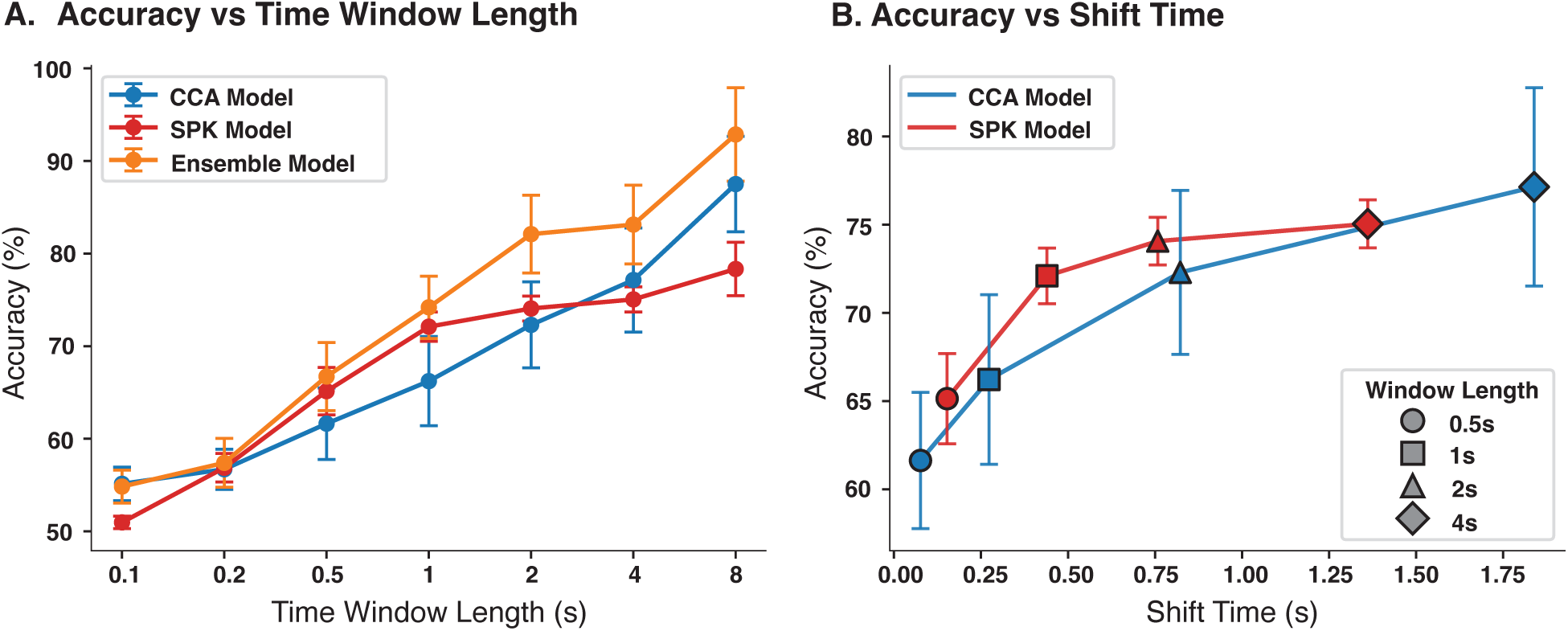
(A) Accuracies of the CCA model, speaker decoding model, and the ensemble model across time windows (0.1–8 s). (B) Attention shift latency for each model. The CCA model responds faster for short windows, while our model shifts faster at longer windows, highlighting a trade-off between speed and accuracy.

To assess how quickly each model responds to a change in a listener’s focus, we simulated attention shifts and measured the detection latency. These trials were created by concatenating neural recordings from two different segments from different speakers, causing the attended speaker to switch midway through the combined segment. Two corresponding stimulus tracks were constructed: one by joining the attended speech of the first sentence with the unattended speech of the second, and the other by reversing this order. To track attention over time, the CCA model used the correlation between the transformed neural and stimulus representations, while our model used the distance between the x-vector of the decoded cluster centroid and the x-vector of the stimulus. The attention shift time was then defined as the duration between the actual moment of the shift and the point at which the model’s tracking metric changed direction to favor the second speaker. The results, presented in Figure 4B, reveal a distinct trade-off. For shorter time windows, our model achieves higher overall accuracy, but the CCA model reacts more quickly to attention shifts. In contrast, for longer windows, the CCA model has a higher accuracy, but our model detects the shifts more rapidly. This shows that our speaker decoding approach exhibits less variability in both accuracy and latency compared to the more time-sensitive CCA-based method.

### 3.4 Model Ablations and Component Analysis

To understand the contribution of each component of our framework, we performed a series of ablation studies (Table 2). We first established performance bounds by simulating a perfect brain predictor (100% accuracy) and a perfect speech extractor. We also established a lower bound using a random cluster prediction, which performed worse than the unprocessed speech mixture. Our experiments confirmed that our design choices were critical. Training a speech extractor on non-quantized x-vectors (xvect-∞) performed worse than using the cluster centroids, validating our clustering approach. Furthermore, we found that our model’s performance is dependent on the strength of its components. A brain predictor with a one-layer bidirectional RNN outperformed simpler (linear, unidirectional) or more complex (2-layer) variants. Similarly, using a stronger speech extractor (Mamba-TasNet L) further improved the results, demonstrating a clear path for future enhancements.

**Table 2:** Upper and lower bounds of our model (xvect-8) and ablations on different model choices.

| Configuration | Pred Acc% | SNR | SI-SDR | PESQ | WER% | SIM% |
| --- | --- | --- | --- | --- | --- | --- |
| *Upper/Lower Bounds* |  |  |  |  |  |  |
| perfect prediction | 100.0 | 13.4 | 12.2 | 2.7 | 9.8 | 95.3 |
| perfect extraction | 78.4 | 84.9 | 82.0 | 4.3 | 13.0 | 98.2 |
| random prediction | 14.0 | 5.0 | -13.5 | 1.8 | 67.6 | 78.2 |
| xvect-∞ extractor | 78.4 | 10.5 | 6.2 | 2.4 | 30.8 | 92.6 |
| *Ablations on Models* |  |  |  |  |  |  |
| linear | 72.0 | 11.8 | 7.3 | 2.5 | 20.8 | 93.4 |
| unidirectional RNN | 77.2 | 12.6 | 10.2 | 2.6 | 15.2 | 94.6 |
| 2-layer RNN | 77.2 | 12.5 | 10.2 | 2.6 | 17.0 | 94.1 |
| Sepformer (Subakan et al., 2021) | 78.4 | 12.0 | 9.8 | 2.5 | 21.0 | 94.0 |
| Mamba-TasNet (L) (Jiang et al., 2024) | 78.4 | 13.5 | 11.2 | 2.8 | 13.5 | 95.4 |

### 3.5 Impact of Speaker Features and Cluster Number

Finally, we investigated how the choice of speaker features and the number of clusters (*K*) influenced performance (Figure 5). We tested features encoding speaker identity (x-vector), speaking style (StyleTTS 2), and general acoustics (WavLM), as well as low-level features like pitch. With *K* = 8, x-vectors yielded the best speech extraction quality and the highest brain prediction accuracy, significantly outperforming all other feature types. We then varied the number of clusters from *K* = 2 to *K* = 32 using x-vectors. As expected, prediction accuracy decreased as *K* increased, since the classification task becomes more difficult. However, the overall extraction performance peaked at *K* = 8. This is because with too few clusters (e.g. *K* = 2), the attended and unattended speakers are more often assigned the same label, making separation impossible. With more clusters (*K* ≥ 8), the extractor can still function correctly even with a prediction error if the predicted cluster is acoustically closer to the attended speaker than the unattended one. We also noted that the performance for *K* = 2 was nearly identical to a gender-based classifier, as the two clusters corresponded almost perfectly to male and female speakers.

**Figure 5:**
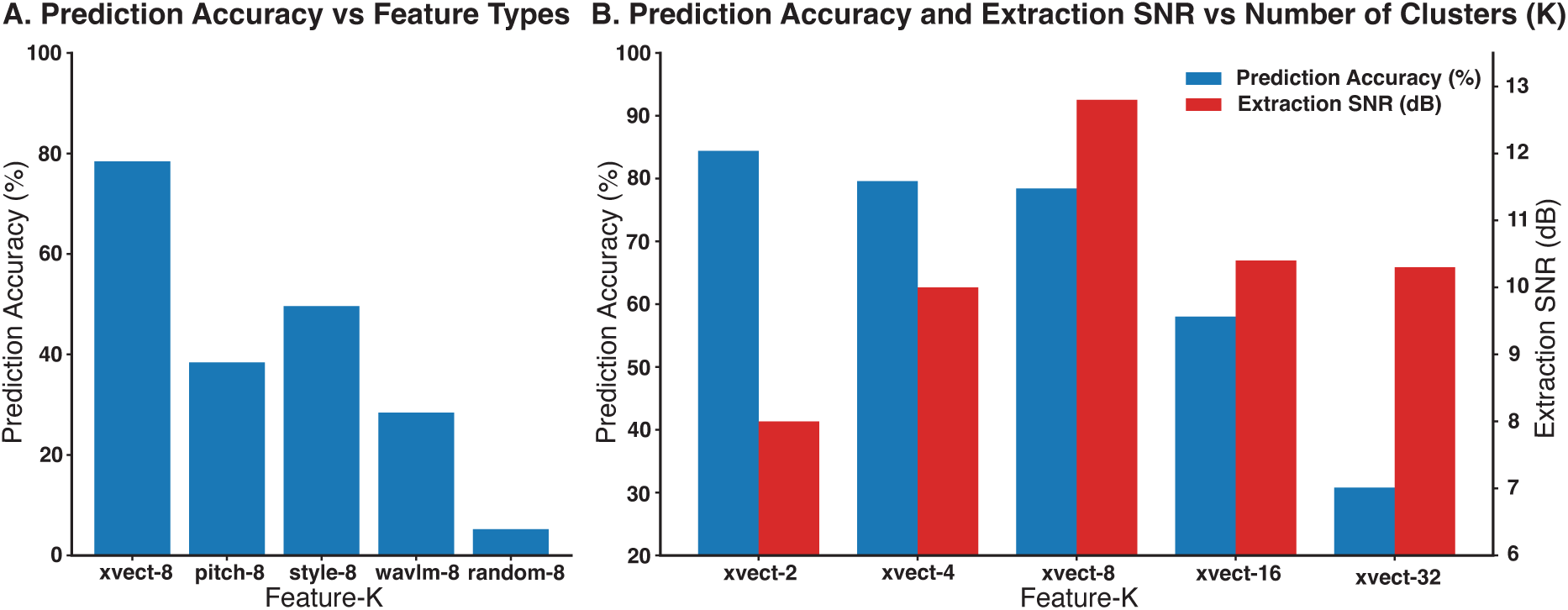
Effect of speaker feature type and cluster number on performance. (A) With *K=8*, X-vectors achieved the highest brain-prediction accuracy and best speech-extraction quality compared to other features. (B) Varying the number of clusters showed that while larger *K* values reduced classification accuracy, extraction performance peaked at *K=8*.

## 4 Discussion

This work introduces a fundamental shift in Auditory Attention Decoding (AAD), moving from reliance on the temporal dynamics of speech to the utilization of time-invariant speaker identity. We demonstrate that the attended speaker is not only tracked temporally but is also robustly encoded in distinct spatial patterns of cortical activity. By leveraging these neural “fingerprints”, our modular system effectively decodes attention and enhances the target voice, achieving performance superior to conventional methods, particularly in challenging short time windows. Furthermore, we show that this spatial mechanism operates in parallel with established temporal processes; a composite model integrating both strategies yields the highest overall performance, highlighting the complementary nature of identifying *who* is speaking versus *when* they are speaking.

Our findings establish a spatial mechanism for attentional selection that operates in parallel with well-documented temporal mechanisms, such as neural phase-locking to the speech envelope (Di Liberto et al., 2022). This represents a shift in the approach to AAD. We show that attention can be decoded by identifying which neural populations are active, rather than how their activity fluctuates over time in correspondence with the stimulus. This spatial approach provides a distinct advantage in temporal resolution; while correlation-based methods require time to accumulate stable evidence (Ceolini et al., 2020; Cheveigné et al., 2018; Han et al., 2019; O’Sullivan et al., 2015), a neural pattern corresponding to a speaker’s identity can, in principle, be identified from a much shorter snapshot of brain activity. This explains our model’s strong performance in time windows of just one to two seconds.

The ability to resolve these fine-grained spatial patterns is critically dependent on the high signal quality and spatial resolution of iEEG (Muller et al., 2016). This presents both a key strength and a primary limitation of our study. The use of invasive recordings provided the necessary spatial resolution to uncover these speaker-specific neural ensembles, which is likely why this mechanism has not been identified previously using lower-resolution, non-invasive methods like scalp EEG. While this reliance on iEEG currently limits the direct clinical translation of our system, it provides an invaluable proof-of-concept and a clear blueprint for what future high-density, non-invasive sensors should aim to capture.

An implication of this work is the potential to decode attentional targets that are not strictly time-locked to stimulus features. Previous AAD systems were fundamentally constrained to decoding features that could be temporally correlated with the neural signal (e.g., envelopes, spectrograms). This limited them to tracking attention to an ongoing stimulus but precluded the ability to decode a listener’s higher-level, sustained intention. Our approach, by decoding a static, categorical feature, opens the door to a new class of neuro-steered technologies. For instance, the same principle could be used to differentiate attention between distinct auditory objects like speech and music, which are known to engage different cortical regions (Norman-Haignere et al., 2015). Looking further, one can envision a system that decodes an abstract intention that is defined not by immediate acoustic properties but by a stable pattern of brain activity. This would represent a major leap from reactive, stimulus-driven BCIs to proactive, intention-driven interfaces (Jiang et al., 2025).

In conclusion, this study demonstrates the viability of a new class of AAD systems based on decoding time-invariant speaker identity from spatial patterns of neural activity. This approach complements existing temporal methods and offers unique advantages in robustness and speed. By shifting the focus from “how” the brain responds over time to “what” populations in the brain are engaged, we not only improve decoding performance but also open a promising new avenue for creating technologies that can interpret and act upon higher-level human intent.

## Data and Code Availability

The data that support the findings of this study are available upon request from N.M. The code for feature extraction, attention decoding, and speech extraction pipelines is available at the following GitHub repository: https://github.com/susameddin/Speaker-Identity-AAD. The code for extracting the x-vector features is available at https://huggingface.co/speechbrain/spkrec-xvect-voxceleb (Snyder et al., 2018). The code for extracting the StyleTTS 2 features is available at https://github.com/yl4579/StyleTTS2 (Li et al., 2024). The code for extracting the WavLM features is available at https://huggingface.co/microsoft/wavlm-base (Chen et al., 2022).

## Author Contributions

S.S.D., X.J., and N.M conceived the project, designed the experiments, evaluated the results, and wrote the manuscript. S.S.D. and X.J. conducted the analysis and model development with support from V.C. and N.M. Surgeries and data acquisition were performed by S.B., A.M., C.S., G.M.M., D.F., and A.F. The project was supervised by N.M. All authors commented on the manuscript.

## Acknowledgment

This work was supported by the National Institutes of Health (NIH) - National Institute on Deafness and Other Communication Disorders (NIDCD) (DC014279) and a grant from Marie-Josee and Henry R. Kravis.

## References

Blei, D.M., Ng, A.Y., Jordan, M.I., 2003. Latent dirichlet allocation. J Mach Learn Res 3, 993–1022.

Breiman, L., 2001. Random Forests. Mach. Learn. 45, 5–32. 10.1023/A:1010933404324

Ceolini, E., Hjortkjær, J., Wong, D.D.E., O’Sullivan, J., Raghavan, V.S., Herrero, J., Mehta, A.D., Liu, S.-C., Mesgarani, N., 2020. Brain-informed speech separation (BISS) for enhancement of target speaker in multitalker speech perception. NeuroImage 223, 117282. 10.1016/j.neuroimage.2020.117282

Chen, S., Wang, C., Chen, Zhengyang, Wu, Y., Liu, S., Chen, Zhuo, Li, J., Kanda, N., Yoshioka, T., Xiao, X., others, 2022. Wavlm: Large-scale self-supervised pre-training for full stack speech processing. IEEE J. Sel. Top. Signal Process. 16, 1505–1518.

Cherry, E.C., 1953. Some experiments on the recognition of speech, with one and with two ears. J. Acoust. Soc. Am. 25, 975–979.

Cheveigné, A. de, Wong, D.D.E., Liberto, G.M.D., Hjortkjær, J., Slaney, M., Lalor, E., 2018. Decoding the auditory brain with canonical component analysis. NeuroImage 172, 206–216. 10.1016/j.neuroimage.2018.01.033

Ciccarelli, G., Smalt, C., Quatieri, T., Brandstein, M., Calamia, P., Haro, S., Nolan, M., Perricone, J., Mesgarani, N., O’Sullivan, J., 2023. End-to-end deep neural network for auditory attention decoding. US11630513B2.

de Cheveigné, A., Wong, D.D.E., Di Liberto, G.M., Hjortkjær, J., Slaney, M., Lalor, E., 2018. Decoding the auditory brain with canonical component analysis. NeuroImage 172, 206–216.

Di Liberto, G.M., Hjortkjær, J., Mesgarani, N., 2022. Editorial: Neural Tracking: Closing the Gap Between Neurophysiology and Translational Medicine. Front. Neurosci. 16. 10.3389/fnins.2022.872600

Ding, N., Simon, J.Z., 2012. Emergence of neural encoding of auditory objects while listening to competing speakers. Proc. Natl. Acad. Sci. 109, 11854–11859.

Formisano, E., De Martino, F., Bonte, M., Goebel, R., 2008. “Who” Is Saying “What”? Brain-Based Decoding of Human Voice and Speech. Science 322, 970–970.

Geirnaert, S., Vandecappelle, S., Alickovic, E., de Cheveigne, A., Lalor, E., Meyer, B.T., Miran, S., Francart, T., Bertrand, A., 2021. Electroencephalography-Based Auditory Attention Decoding: Toward Neurosteered Hearing Devices. IEEE Signal Process. Mag. 38, 89–102. 10.1109/MSP.2021.3075932

Gu, A., Dao, T., 2023.Mamba: Linear-time sequence modeling with selective state spaces. ArXiv Prepr. ArXiv231200752.

Han, C., O’Sullivan, J., Luo, Y., Herrero, J., Mehta, A.D., Mesgarani, N., 2019. Speaker-independent auditory attention decoding without access to clean speech sources. Sci. Adv. 5, eaav6134–eaav6134.

Hochreiter, S., Schmidhuber, J., 1997. Long Short-Term Memory. Neural Comput. 9, 1735–1780. 10.1162/neco.1997.9.8.1735

Jiang, X., Dindar, S.S., Choudhari, V., Bickel, S., Mehta, A., McKhann, G.M., Friedman, D., Flinker, A., Mesgarani, N., 2025.AAD-LLM: Neural Attention-Driven Auditory Scene Understanding.

Jiang, X., Li, Y.A., Florea, A.N., Han, C., Mesgarani, N., 2024.Speech Slytherin: Examining the Performance and Efficiency of Mamba for Speech Separation, Recognition, and Synthesis. ArXiv Prepr. ArXiv240709732.

Kingma, D.P., Ba, J., 2014. Adam: A Method for Stochastic Optimization. CoRR abs/1412.6980.

Le Roux, J., Wisdom, S., Erdogan, H., Hershey, J.R., 2019. SDR–half-baked or well done?, in: ICASSP 2019-2019 IEEE International Conference on Acoustics, Speech and Signal Processing (ICASSP). IEEE, pp. 626–630.

Li, Y.A., Han, C., Raghavan, V., Mischler, G., Mesgarani, N., 2024.Styletts 2: Towards human-level text-to-speech through style diffusion and adversarial training with large speech language models. Adv. Neural Inf. Process. Syst. 36.

Maaten, L. van der, Hinton, G., 2008. Visualizing Data using t-SNE. J. Mach. Learn. Res. 9, 2579–2605.

Mesgarani, N., Chang, E.F., 2012. Selective cortical representation of attended speaker in multi-talker speech perception. Nature 485, 233–236.

Mesgarani, N., David, S.V., Fritz, J.B., Shamma, S.A., 2008. Phoneme representation and classification in primary auditory cortex. J. Acoust. Soc. Am. 123, 899–909. 10.1121/1.2816572

Muller, L., Hamilton, L.S., Edwards, E., Bouchard, K.E., Chang, E.F., 2016. Spatial resolution dependence on spectral frequency in human speech cortex electrocorticography. J. Neural Eng. 13, 56013–56013.

Norman-Haignere, S., Kanwisher, N.G., McDermott, J.H., 2015. Distinct cortical pathways for music and speech revealed by hypothesis-free voxel decomposition. Neuron 88, 1281–1296.

O’Sullivan, J., Chen, Z., Herrero, J., McKhann, G.M., Sheth, S.A., Mehta, A.D., Mesgarani, N., 2017. Neural decoding of attentional selection in multi-speaker environments without access to clean sources. J. Neural Eng. 14. 10.1088/1741-2552/aa7ab4

O’Sullivan, J., Herrero, J., Smith, E., Schevon, C., McKhann, G.M., Sheth, S.A., Mehta, A.D., Mesgarani, N., 2019. Hierarchical Encoding of Attended Auditory Objects in Multi-talker Speech Perception. Neuron 104. 10.1016/j.neuron.2019.09.007

O’Sullivan, J.A., Power, A.J., Mesgarani, N., Rajaram, S., Foxe, J.J., Shinn-Cunningham, B.G., Slaney, M., Shamma, S.A., Lalor, E.C., 2015. Attentional Selection in a Cocktail Party Environment Can Be Decoded from Single-Trial EEG. Cereb. Cortex 25, 1697–1706.

Patel, P., Heijden, K. van der, Bickel, S., Herrero, J.L., Mehta, A.D., Mesgarani, N., 2022. Interaction of bottom-up and top-down neural mechanisms in spatial multi-talker speech perception. Curr. Biol. 32, 3971–3986.e4. 10.1016/j.cub.2022.07.047

Patel, P., Long, L.K., Herrero, J.L., Mehta, A.D., Mesgarani, N., 2018. Joint Representation of Spatial and Phonetic Features in the Human Core Auditory Cortex. Cell Rep. 24, 2051–2062.

Pedregosa, F., Varoquaux, G., Gramfort, A., Michel, V., Thirion, B., Grisel, O., Blondel, M., Prettenhofer, P., Weiss, R., Dubourg, V., Vanderplas, J., Passos, A., Cournapeau, D., Brucher, M., Perrot, M., Duchesnay, E., 2011. Scikit-learn: Machine Learning in Python. J. Mach. Learn. Res. 12, 2825–2830.

Perez, E., Strub, F., Vries, H. de, Dumoulin, V., Courville, A., 2017. FiLM: Visual Reasoning with a General Conditioning Layer.

Radford, A., Kim, J.W., Xu, T., Brockman, G., McLeavey, C., Sutskever, I., 2023. Robust speech recognition via large-scale weak supervision, in: International Conference on Machine Learning. PMLR, pp. 28492–28518.

Rix, A.W., Beerends, J.G., Hollier, M.P., Hekstra, A.P., 2001. Perceptual evaluation of speech quality (PESQ)-a new method for speech quality assessment of telephone networks and codecs, in: 2001 IEEE International Conference on Acoustics, Speech, and Signal Processing. Proceedings (Cat. No.01CH37221). pp. 749–752 vol.2. 10.1109/ICASSP.2001.941023

Shamma, S.A., Elhilali, M., Micheyl, C., 2011. Temporal coherence and attention in auditory scene analysis. Trends Neurosci. 34, 114–123. 10.1016/j.tins.2010.11.002

Snyder, D., Garcia-Romero, D., Sell, G., Povey, D., Khudanpur, S., 2018. X-Vectors: Robust DNN Embeddings for Speaker Recognition, in: 2018 IEEE International Conference on Acoustics, Speech and Signal Processing (ICASSP). pp. 5329–5333.

Tanveer, M.A., Skoglund, M.A., Bernhardsson, B., Alickovic, E., 2024. Deep learning-based auditory attention decoding in listeners with hearing impairment*. J. Neural Eng. 21, 036022. 10.1088/1741-2552/ad49d7

Theunissen, F.E., Sen, K., Doupe, A.J., 2000. Spectral-temporal receptive fields of nonlinear auditory neurons obtained using natural sounds. J Neurosci 20, 2315–2331.

van der Heijden, K., Patel, P., Bickel, S., Herrero, J.L., Mehta, A.D., Mesgarani, N., 2025. Joint population coding and temporal coherence link an attended talker’s voice and location features in naturalistic multi-talker scenes. J. Neurosci. 45.

Walker, K.M.M., Bizley, J.K., King, A.J., Schnupp, J.W.H., 2011. Multiplexed and Robust Representations of Sound Features in Auditory Cortex. J. Neurosci. 31, 14565–14576. 10.1523/JNEUROSCI.2074-11.2011

Zen, H., Dang, V., Clark, R., Zhang, Y., Weiss, R.J., Jia, Y., Chen, Z., Wu, Y., 2019. LibriTTS: A Corpus Derived from LibriSpeech for Text-to-Speech, in: Interspeech 2019. pp. 1526–1530. 10.21437/Interspeech.2019-2441

Zhu, L., Liao, B., Zhang, Q., Wang, Xinlong, Liu, W., Wang, Xinggang, 2024. Vision mamba: Efficient visual representation learning with bidirectional state space model. ArXiv Prepr. ArXiv240109417.

